# Temporal dynamics of macrophages transcriptional profiles during zebrafish wound healing

**DOI:** 10.1101/2025.09.12.675595

**Authors:** Christina Begon-Pescia, Resul Özbilgiç, Stéphanie Boireau, Anaïs Louis, Simon Georges, Laurent Manchon, Mai Nguyen-Chi

**Affiliations:** LPHI, Université de Montpellier, CNRS, INSERM, Montpellier, France; Montpellier Ressources Imagerie, Biocampus, CNRS, INSERM, Université Montpellier France; MGX-Montpellier GenomiX, Univ. Montpellier, CNRS, INSERM, Montpellier France; Independent researcher

**Author notes:** Corresponding author(s) (M. Nguyen-Chi).

**Keywords:** zebrafish, transcriptomic profile, macrophages, polarization dynamic, inflammation, wound healing

## Abstract

Macrophages participate to wound healing by contributing to host defense and orchestrating inflammation and repair. They adapt dynamically to the wound microenvironment by adopting diverse polarization states, which are determinant of wound outcome, influencing whether healing is successful or becomes chronic. The zebrafish embryo, widely used for live imaging of immune responses, is a powerful model to study macrophage behavior after injury. However, the transcriptional landscape of polarized macrophages in this model during wound healing remains insufficiently characterized. Here, we employ bulk RNA sequencing to characterize macrophage transcriptional programs following tail fin wounding a robust model for studying sterile inflammation. Our findings reveal that zebrafish macrophages undergo large transcriptomic changes along different wound healing phases, particularly between 2 and 5 hours post-amputation, suggesting a fast reprogramming leading to different functional states. We further show that, at 2h, macrophages acquire a pro-inflammatory profile with a gene signature closed to M1 signature. At 5h, macrophages express genes involved in immunoregulation and healing associated with shutoff of pro-inflammatory pathways and the activation of glucose and glycogen metabolism. Finally, we show that macrophage reprogramming becomes deeply attenuated by 29h. Our findings provide a foundation for understanding macrophage polarization in zebrafish, revealing underpinning molecular mechanisms, including both specific and evolutionarily conserved pathways with a potential impact on translational medicine.

## 1. Introduction

Wound healing is a complex process that involves multiple cell populations interacting in a coordinated manner during successive phases: inflammation, proliferation, re-epithelialization, and remodeling^1^. The restoration of skin integrity is essential not only for protecting the host from pathogens but also for re-establishing barrier function. Despite extensive research, effective therapeutic strategies that accelerate tissue repair while reducing scarring remain limited. A deeper understanding of the roles of various cell populations at play in wound repair is therefore essential to elucidate the mechanisms underlying normal wound closure and to prevent the development of pathological wounds.

Macrophages are among the first responders recruited to the injury site, where they play pivotal roles in both host defense and tissue repair. As key regulators of wound healing, macrophages modulate the onset and resolution of inflammation and orchestrate several processes integral to tissue repair^2^. Numerous studies have demonstrated that macrophages are indispensable for proper wound healing and regeneration across a range of species, from mammals^3,4^ to non-mammalian models^5,6,7,8,9^.

As the most plastic cells within the hematopoietic system, macrophages adapt to microenvironmental cues through a process known as polarization. This process, driven by specific gene expression programs, enables macrophages to acquire distinct functional phenotypes. *In vitro*, macrophages were classified into two major groups: the pro-inflammatory M1 (classically activated) and the anti-inflammatory M2 (alternatively activated) states^10,11^. At the onset of wound healing, macrophages predominantly adopt an M1 phenotype characterized by the secretion of pro-inflammatory cytokines such as tumor necrosis factor (TNF) and interleukin-6 (IL-6). These M1 macrophages are critical for host defense and the clearance of cellular debris, and they contribute to the stimulation of cell proliferation in the damaged tissue^12^. As healing progresses, a phenotypic shift occurs toward an M2 state, evidenced by increased expression of markers such as MRC1. M2 macrophages secrete growth factors most notably transforming growth factor-β1 (TGF-β1) which promote stromal cell proliferation, survival, and migration. In addition, these cells facilitate the resolution of inflammation and mediate reparative processes including angiogenesis, granulation tissue formation, re-epithelialization, and extracellular matrix remodeling through the release of matrix metalloproteases^12,13^. However, this classification has been challenged by *in vivo* studies^14^. In addition single-cell RNA sequencing studies have revealed that macrophage phenotypes during wound healing are more heterogeneous than previously appreciated^15,16^. Importantly, the precise temporal regulation of macrophage polarization is essential for normal healing, and its impairment leads to delay of tissue repair and regeneration in various animal models^17,6,8^. Despite these advances, the full spectrum of macrophage states in wounds remains incompletely characterized, and it is essential to associate each state with its corresponding functional behavior.

Zebrafish (*Danio rerio*) is an excellent model for studying wound healing mechanisms due to its remarkable regenerative abilities and its transparent embryos, which permit real-time observation of cell populations^18,19,20^. The strong genetic resemblance to humans further underscores its relevance for biomedical research^21^. Recently, there has been increasing focus on the role of the immune system, especially macrophages, in wound repair and regeneration. Zebrafish macrophages, like their mammalian counterparts, contribute to both antimicrobial and pro-inflammatory responses^22,23,24,20^ as well as to tissue repair^6,8,9,25^. Macrophages are crucial in wound healing, including in the caudal fin of both adult and larval zebrafish^8,9^. During the early phases of inflammation, zebrafish macrophages adopt an M1-like phenotype, marked by cytokine production such as *tnfa.a*, *tnfa.b*, *il1b*, and *il6*. In the later stages, macrophages transition to an M2-like, characterized by the expression of *tgf-b1* and *ccr2* and *cxcr4* ^26,27,8,24^. Another marker associated with wound macrophage subsets in later stages is *arg2*, which has been suggested to mediate anti-inflammatory functions^28^. Yet, the detailed molecular programs, as well as the timing of macrophage activation during tissue injury in zebrafish are poorly defined. In addition, markers to track distinct wound-macrophage subsets are still lacking in this specie.

Here, we used transcriptomic profiling of macrophages to identify the changes in gene expression that control the molecular mechanisms underlying the macrophage polarization during wound healing in zebrafish. Using a transgenic zebrafish macrophage reporter line, we isolated macrophages by FACS cell sorting at different time points after caudal fin fold injury and analyzed their transcriptome by bulk RNA-sequencing. We found that the gene expression patterns of macrophages are highly dynamic during the healing process. We identified transcriptomic programs and biological pathways that distinguish the acute phase of inflammation, transition to resolution and remodeling phase.

## 2. Methods

### Ethics statement

All animal experiments were conducted by well-trained and authorized staff in compliance with the European Union guidelines for handling of laboratory animals (http://ec.europa.eu/environment/chemicals/lab_animals/index_en.htm) and the ARRIVE guidelines. Breeding and maintenance of adult fish were performed at the ZEFIX-LPHI (CNRS, University of Montpellier, Montpellier, France; license number CEEA-LR-B34–172–37). Developmental stages used were 3 dpf and 4 dpf. When required, specimens were euthanized using an anesthetic overdose of buffered tricaine (500 mg/L).

### Fish husbandry

Zebrafish (*Danio rerio*) maintenance and husbandry were performed at the fish facility of the University of Montpellier as describes^29^. Males and females were kept in 3.5-L polycarbonate tank (maximum 22 fish per tank) connected to a recirculating system (Tecniplast), in the following conditions: 4 ‰ salinity / 400 conductivity, temperature of 27.5°C and a 12:12-h light: dark cycle. The fish were fed twice per day. Embryos were obtained from pairs of adult fishes by natural spawning at 28°C in tank water. Embryos and larvae were staged according to Kimmel et al.^30^ in zebrafish medium in 10-cm petri dishes. The zebrafish embryo medium was changed every day. Strains used were transgenic lines: *Tg (mfap4:mCherry-F)* ump6Tg, referred as *Tg (mfap4:mCherry-F)*^31^ to specific label the macrophages with a farnesylated mCherry protein and AB strain, referred as WT, was used as a control for FACS.

### Zebrafish caudal fin fold amputation and cell dissociation

Larvae were anaesthetized in zebrafish medium supplemented with 200 μg / ml tricaine (ethyl 3-aminobenzoate methanesulfonate, MS-222 sigma # A5040) before any manipulation. Caudal fin fold amputation was performed at 3 dpf for all experiments as previously described^8^. Briefly, fin folds were transected with a sterile scalpel, posterior to muscle and notochord. Larvae were transferred in their medium at 28°C for 2, 5 and 29 hours and they correspond to 2 hours post-amputation (hpA), 5 hpA and 29 hpA group. 360 *Tg (mfap4:mCherry-F)* larvae were either amputated (CUT) or not amputated as a control (UNcut), then crushed on cell strainer 70-μm mesh in cold working buffer (0.9X DPBS with 2% FBSi and 2 mM EDTA). Isolated cells were washed, filtered through a cell strainer 40-μm mesh and transferred into 5 ml round bottom polystyrene FACS tubes as previously describe^29^. A total of four replicates for each condition (CUT ou UNcut) within each group (2 hpA, 5 hpA and 29 hpA) were conducted.

### FACS cell sorting and mRNA isolation from sorted macrophage cells

Following cell dissociation from amputated and non-amputated larvae, mCherry-F positive cells were sorted using a FACSAria^TM^ IIu (BD Biosciences) as previously describe^29^. To minimize RNA degradation and material loss, the mCherry-F positive and mCherry-F negative sorted cells were separately collected directly into RNase/DNase-free 1.5 ml microfuge tubes containing with 100 μl RLT/β-ME Qiagen’s lysis buffer and kept on ice for immediate lysis. A set of 24 total RNA extractions from the mCherry-F positive sorted macrophages were performed using RNeasy^®^ Micro Kit from Qiagen according to manufacturer’s instructions. Between 30 000 and 97 000 mCherry-F positive cells were sorted, yielding a total RNA amount ranging from 5 to 50 ng per sample. All RNA integrity number (RIN) obtained were above 7.

### Illumina sequencing and Differential Gene Expression analysis (DEGs) on FACS-sorted macrophages

The sequencing data generated by this study have deposited in the NCBI Gene Expression Omnibus (GEO) under the accession number GSE234476. We generated the RNA-Seq libraries with the Illumina DNA prep kit from 5 to 10 ng input RNA. The cDNA synthesis and amplification were performed using the SMART-Seq^®^ V4 Ultra Low kit (Takara Bio). The libraries have been sequenced using a 100-nucleotides single-read SP flow cell lane using the 2-channel Sequence By Synthesis (SBS) method. The libraries were validated by quantification of the DNA concentration and fragment length using the Fragment Analyzer (Standard Sensitivity NGS kit, Agilent Technologies, Santa Clara, CA, USA) and qPCR (Roche LightCycler®480). FeatureCounts (v2.0.3) was used to count the number of reads for each sample, which ranged between 30 and 48 million at 2 hpA, between 35 and 51 million at 5 hpA and between 35 and 45 million at 29 hpA. RNA-sequencing raw reads were then aligned to the *Danio rerio* reference genome (Ensembl, GRCz11) using TopHat2 software^32^. The “raw read count” data sets were analyzed using the Bioconductor package DESeq2 (v1.38.2)^33,34^ to identify the differentially expressed genes (DEGs) between CUT and UNcut conditions at different times post-amputation (2, 5 hpA) with cut off Log_2_FC |1| and 29 hpA (cut off Log_2_FC |0|) and adjusted *p* value ≤ 0.05 (according to the FDR method from Benjamini-Hochberg^35^ as the significance cut-off to identify DEGs).

### RNA sequencing bioinformatics data analysis, GO and pathway analysis

The transcriptome profiling of macrophages during wound healing was analyzed using Bioconductor^34^ in R (v4.1.2) with the following packages: ggvenn, ggplot2, EnhancedVolcano, pheatmap (R Core Team: https://www.R-project.org/). Venn diagrams were generated using ggvenn to illustrate the overlap between different sets of DEGs and identify common genes across the time course. A scatter-plot was created using ggplot2 to depict the fold-change correlation between all common genes identified in the Venn diagram. To visualize gene expression changes at different time points, we constructed Volcano plots, using EnhancedVolcano. Additionally, hierarchical clustering heatmaps were generated using pheatmap to display the expression patterns of the top 50 most up-regulated and down-regulated genes, comparing CUT conditions to UNcut conditions within each group. Finally, to functionally categorize the differentially expressed genes, we performed Gene Ontology (GO) analysis to determine their biological roles based on the biological process, the molecular function enrichment and the Kyoto Encyclopedia of Genes and Genomes (KEGG) pathway annotations. This analysis was conducted using ShinyGO v0.77^36^ and SRplot^37^.

### RNA-Sequencing validation by quantitative Real Time PCR assays and statistical analysis

Validation of RNA-sequencing data was assessed using quantitative Real Time PCR (RT-qPCR). We selected several differentially expressed genes, including both up-regulated and down-regulated genes for each group. Primers were designed with Primer 3 Plus and Primer-BLAST online tool (https://www.primer3plus.com/index.html and https://www.ncbi.nlm.nih.gov/tools/primer-blast/). Zebrafish *ef1a* (*eukaryotic translation elongation factor 1 alpha 1, like 1*) was used as the reference gene for normalization to determine the relative gene expression levels. Primer sequences and amplicon size for the following zebrafish genes was listed in Table 1: *il-1b, cxcl8 (il-8), tnfa.a, il6, cxcl18b, cxcr4b, tlr8a, tlr2, sox4b, s1pr4, timp2b, arg2, f5, mmp9, c1qb, dicp1.1, tnfa.b, p2rx7, sulf2b*. Each primer set was tested for amplification efficiency before use. For cDNA synthesis, 10 ng of total RNA was reverse-transcribed using Oligo (dT)_18_ primers and M-MLV reverse transcriptase (Invitrogen^TM^ # 28025-013) following the manufacturer’s recommendations. RT-qPCR was performed using a LightCycler® 480 (Roche) with SYBER Green (Meridian Bioscience^®^ SensIFAST^TM^ SYBR^®^ No-ROX Mix # BIO-98050), according to the manufacturer’s protocol. All experiments were performed in both biological and technical replicates, ranging from 3 to 7 biological replicates per time point. Samples were run in triplicate and data were analyzed using LightCycler^®^ 480 software (version 1.5.1). Zebrafish *ef1a* (*eukaryotic translation elongation factor 1 alpha 1, like 1*) was used as the reference housekeeping gene for normalization and the relative gene expression levels were calculated using the 2^-ΔCt^ method (target / reference). The data analysis was performed using GraphPad Prism 8 software (version 8.3.0). Results were visualized as scatter plots with individual values, bar means and standard error mean (SEM). Statistical significance was evaluated using Mann-Whitney U tests (unpaired, non-parametric), with a significance threshold set at p-value < 0.05.

### Fold change correlation between RNA-Sequencing and quantitative Real Time PCR data

To evaluate the correlation between RNA-sequencing data and RT-qPCR results, we assessed a Spearman’s rho correlation coefficient of the fold change. Several DEGs per time point were taken randomly from the RNA-Seq datasets. Their fold change was then compared with that of the RT-qPCR results.

### Imaging of living zebrafish larvae

Larvae were anesthetized in 200-μg / ml tricaine (Ethyl 3-aminobenzoate 646 methanesulfonate, MS-222 Sigma), positioned in 35 mm glass-bottom dishes (FluoDish, World Precision Instruments, UK), immobilized in 1% low melting point agarose (Sigma) and covered with 2 ml of embryo water containing tricaine. For imaging, confocal microscopy was performed on Zeiss LSM880 FastAiryscan, using 20X/0.8 objective. The wavelength was 561 nm for excitation. The images were taken in a sequential mode by line. The 3D files generated by multi-scan acquisitions were processed by Image J software. Brightness and contrast were adjusted for maximal visibility.

### Data and code availability

The RNA-sequencing raw data from this study have been submitted and archived to the NCBI Gene Expression Omnibus (https://www.ncbi.nlm.nih.gov/geo/) under sample accession number GSE234476.

## 3. Results

### Comparative analysis of zebrafish macrophage transcriptomes revealed distinct profiles across wound healing

To study wound healing, we used the well-established model of caudal fin fold amputation in 3 dpf zebrafish larvae. In order to accurately determine the acute inflammatory phase, we performed a global analysis of pro-inflammatory marker expression in the whole larvae at various times after wound: 0, 1, 2, 3, 5, 6 and 24 hours post-amputation. The induction kinetics of *il1b, il8, tnfa.b (here named tnfb), nos2a* mRNAs were thus monitored by RT-qPCR in amputated (CUT) versus non-amputated (UNcut) conditions (Figure S1). An increase in *il1b and tnfb* was observed in the amputated condition. Notably, *il1b* and *tnfb* reached their highest expression levels between 1 and 2 hours post-amputation. In contrast, *nos2a and il8* were detected but not significantly modulated (Figure S1). These results indicate that in whole tissue larva, the acute inflammatory phase occurs within the first few hours following injury, with a clear decline by 5–6 hpA, marking the end of acute phase.

Therefore, we decided to analyze the transcriptomic profile of zebrafish macrophages during wound healing at 3 different time points: 2 hpA (corresponding to the acute phase), 5 hpA (time corresponding to the end of acute phase) and 29 hpA (time corresponding to resolution-return to steady state)^38^. To molecularly characterize activated macrophages during this process, we performed a caudal fin fold amputation on macrophage reporter larvae, *Tg (mfap4: mCherry-F)* (Figure S2) and macrophages were sorted by FACS based on their fluorescence at 2, 5 and 29 hpA (Figure S3). Pure populations of macrophages were obtained as previously shown^39^(Figure S4) and global gene expression analysis was performed using bulk RNA-sequencing. All the time points were performed with four biological replicates. To identify differentially expressed genes (DEGs) at each time point, we compared the amputated (CUT) and non-amputated (UNcut) conditions using an adjusted *p*-value ≤ 0.05 as the significance threshold. We found that 1,895 genes were significantly differentially expressed at 2 hpA (Table 2), while 4,216 genes were significantly differentially expressed at 5 hpA (Table 3), and 45 genes at 29 hpA (Table 4). Considering the most modulated genes (cut off Log_2_FC |1|), the volcano plot at 2 hpA, showed 599 genes were up-regulated and 224 down-regulated, while at 5 hpA, 414 genes were up-regulated and 609 down-regulated (Figure 1A, B). At 29 hpA, most DEGs have a Log_2_FC <1 (cutoff Log_2_FC |0|) so, 17 genes were down-regulated and 28 up-regulated (Figure 1C), indicating that macrophages in CUT condition are very similar to that of UNcut condition.

**Figure 1.**
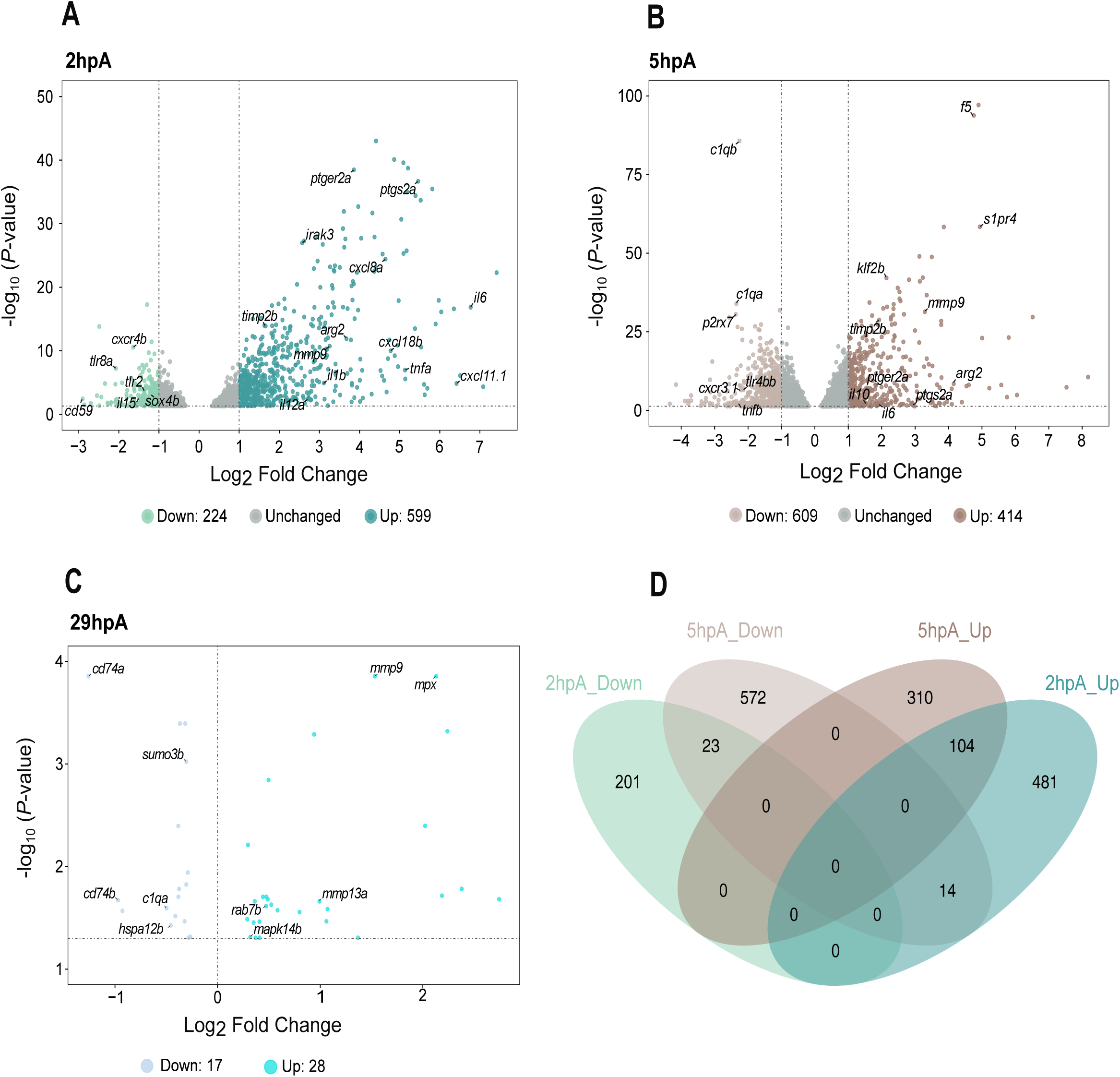
Volcano plots of RNA-sequencing data at 2, 5, and 29 hours post-amputation. Volcano plots illustrating gene expression changes at different time points following injury. Each dot represents a gene. The x-axis shows the log₂ fold change (log₂FC) in gene expression between the amputated (cut) and control (uncut) conditions. The vertical dashed lines mark the fold change threshold (log_2_FC |1|) used for 2 and 5 hpA; no fold change cutoff was applied at 29 hpA. The y-axis shows the -log₁₀ of the Benjamini-Hochberg adjusted p-value (with a significance threshold of p < 0.05), indicated by the horizontal dashed line. Colors denote up-regulated and down-regulated genes. **(A)** At 2 hpA, 599 genes are up-regulated and 224 genes are down-regulated, with the remaining genes unchanged. **(B)** At 5 hpA, 414 genes are up-regulated and 609 genes are down-regulated, with the rest unchanged. **(C)** At 29 hpA, 28 genes are up-regulated and 17 genes are down-regulated. **(D)** The Venn diagram illustrates the number of up-regulated (Up) and down-regulated (Down) DEGs at 2 hours post-amputation (hpA; green) and 5 hpA (brown). Each hemisphere represents the DEGs identified at each time point, with the overlapping region indicating shared genes across both conditions. A total of 823 genes were significantly regulated at 2 hpA, 1,023 at 5 hpA, with 141 DEGs shared between the two time points.

Next, we compared the macrophage transcriptomes at 2 and 5 hpA. Interestingly, the Venn diagram in Figure 1D showed 141 DEGs were shared. However, many DEGs remained specific to each healing stage (682 genes for 2 hpA and 882 genes for 5 hpA)

Collectively, these results show that zebrafish macrophages underwent large transcriptomic changes along different wound healing phases, and particularly between 2 hpA and 5 hpA, suggesting that reprogramming is rather fast and leads to different functional states.

### Gene expression changes observed by RNA sequencing corroborate the results of Real Time PCR

For validating RNA-sequencing data, several DEGs per time points were randomly selected and analyzed by RT-qPCR. For 2 hpA, we studied *cxcr4b, sox4b, tlr8a and tlr2* (down-regulated) and *il-1b, cxcl8a, il-6, tnfa.a* and *cxcl18b* (up-regulated). We found that all the genes that were down-regulated in sequencing data, were also down-regulated in RT-qPCR data, with the exception of *tlr2*, which followed the same down-regulated trend, but was not significant (Figure S5A and Figure 2A). Similarly, all the genes that were up-regulated in sequencing data were also up-regulated in RT-qPCR data (Figure S5B and Figure 2A). The Spearman’s rho correlation coefficient of the log2 fold change was evaluated and showed that results from RNA-sequencing data and RT-qPCR were similar (rho = 0.8, p = 0.014) (Figure 2B).

**Figure 2.**
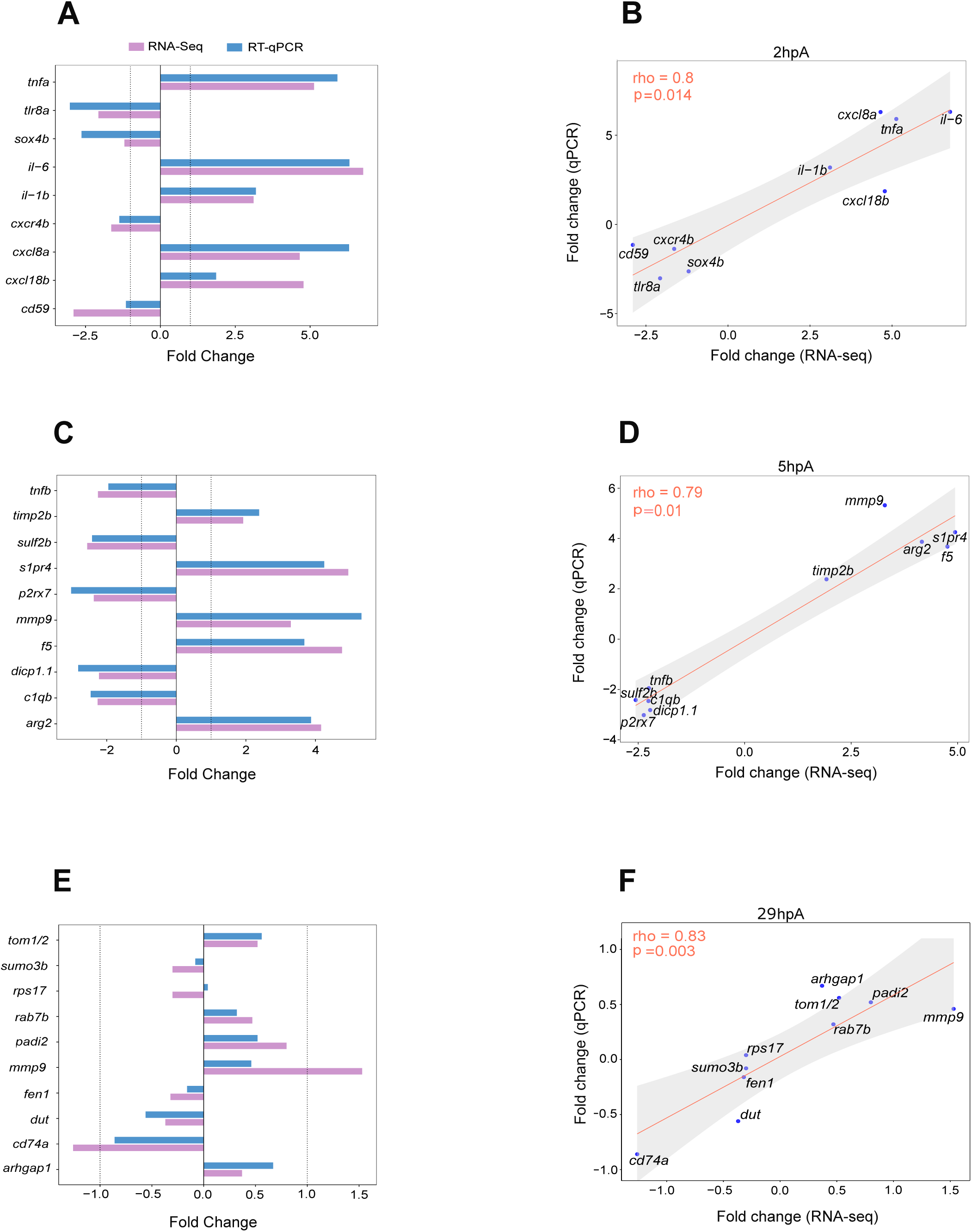
Correlation of fold changes between RT-qPCR and RNA-seq data. Bar plots comparing expression profiles of selected genes at each time point, as measured by RNA-seq (magenta) and RT-qPCR (blue). Gene expression was normalized to the *ef1a* reference gene. Results are presented as mean fold change values **(A, C, E)**. Correlation of gene expression fold changes between RNA-seq and RT-qPCR was assessed using Spearman’s rho correlation coefficient. Several genes were sampled per time point for comparison. **(B, D, F)** Fold change correlation analyses between RNA-seq and RT-qPCR at 2, 5, and 29 hours post-amputation (hpA), respectively. Log₂ fold change values were used for both methods. Spearman’s correlation results: at 2 hpA: rho = 0.8, p = 0.014; at 5 hpA: rho = 0.79, p = 0.01; at 29 hpA: rho = 0.83, p = 0.003).

A comparable analysis was performed at 5 hpA, with randomly selected DEGs: *c1qb, dicp1.1, tnfb, p2rx7* and *sulf2b* (down-regulated genes) and *s1pr4, timp2b, arg2, f5* and *mmp9* (up-regulated genes). Genes that were down-regulated in sequencing data, were also down-regulated in RT-qPCR data, with the exception of *tnfb* which followed the same down-regulated trend, but was not significant (Figure S5C and Figure 2C). Some genes that were up-regulated in sequencing data, were also up-regulated in RT-qPCR data with the exception of *mmp9* and *f5* which followed the same up-regulated trend, but were not significant (Figure S5D and Figure 2C). When compared, the log2 values of the fold change from sequencing data and RT-qPCR data gave a Spearman’s rho, rho = 0.79, p = 0.01 (Figure 2D). All of this suggests an acceptable level of concordance between RNA-sequencing and RT-qPCR results.

At 29 hpA, RT-qPCR analysis with randomly selected DEGs revealed that *cd74a, sumo3b, dut, fen1* and *rps17* showed a down-regulated trend (Figure S5E and Figure 2E). Similarly, *mmp9, padi2, tom1/2, rab7b* and *arhgap1* showed an up-regulated trend (Figure S5F and Figure 2E). In addition, the Spearman’s rho correlation coefficient of the log2 fold change was evaluated and showed that results from RNA-sequencing data and RT-qPCR had an acceptable level of concordance (rho = 0.83, p = 0.003) (Figure 2F).

All of this suggests an overall good level of concordance between RNA-sequencing and RT-qPCR results for 2 hpA, 5 hpA and 29 hpA time points. These data confirm that macrophages modulate their profile during the first steps of wound healing.

### Two-hour-post-amputation macrophages display an acute pro-inflammatory profile

Zebrafish macrophages were previously shown to express pro-inflammatory makers after an immune challenge^27,40^, however the exact molecular program driving M1-like activation during wound healing in this species is incompletely understood. At 2 hpA, we detected pro-inflammatory genes among the top 50 up-regulated genes including *tnfa*, *il6, il11, cxcl18b, cxcl8a* and *cxcl11.1* (Figure 3A and Table 5A).

**Figure 3.**
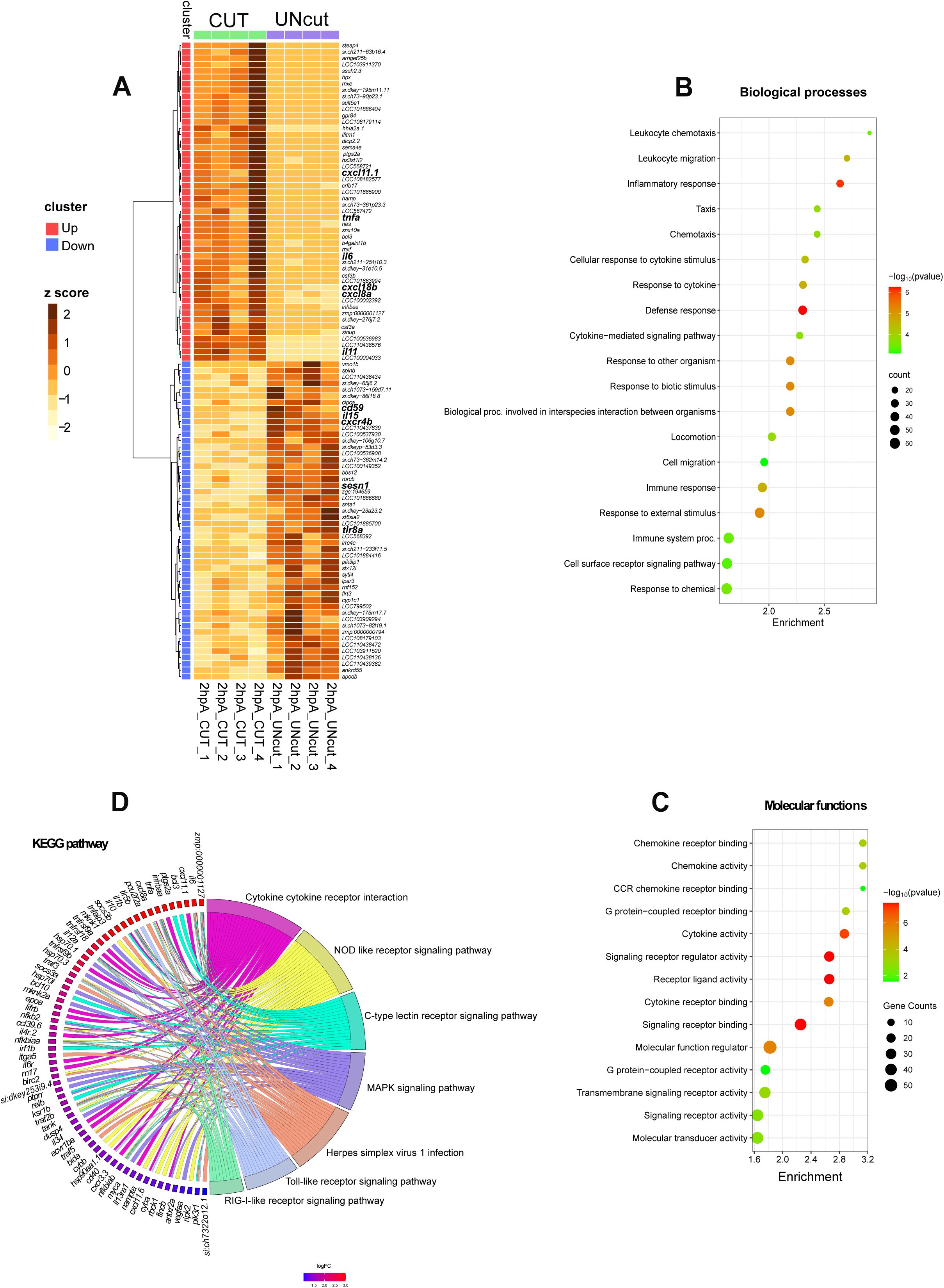
Transcriptome profiling of macrophages at 2 hpA in zebrafish larvae. (**A**) *Heat-map of differentially expressed genes.* The 50 most strongly up- and down-regulated genes are displayed. For each gene, normalized read counts were Z-score–transformed across all samples (columns = individual biological replicates). Colour scale: dark brown = high expression; light yellow = low expression. Clusters enriched in up-regulated genes are marked with red outlines; clusters enriched in down-regulated genes are marked in blue. Samples from amputated larvae (CUT) are labelled with green squares, and control samples (UNcut) with purple squares. Hierarchical clustering was performed with Ward’s minimum-variance method on log₂ fold-change values. **(B, C)** *Bubble plots of Gene Ontology enrichment.* Biological-process (**B**) and molecular-function (**C**). Terms enriched among the up-regulated DEGs are shown. Dot size represents the number of genes in each term; dot colour encodes significance (FDR): red = higher significance (lower *p*-value), green = lower significance (higher *p*-value). The x-axis shows fold enrichment (gene ratio); the y-axis lists enriched GO terms. **(D)** *GO chord plot of Kyoto Encyclopedia of Genes and Genomes (KEGG)-associated terms.* Genes are connected to their associated pathways by ribbons. Genes are ordered by log₂ fold change (red = strong up-regulation; blue = strong down-regulation).

Up-regulated genes at 2 hpA were classified according the gene ontology (GO) terms and the KEGG pathways. GO terms related to biological processes such as “inflammatory response”, “defense response” and “response to cytokine” were enriched from up-regulated genes (Figure 3B). In the molecular functions, the enriched GO terms were: “cytokine activity” (*il-1b, il-12bb*, *il-10*, *il-6*, *il34*, *tnfa*), “molecular function regulator” (*socs1a*, *socs3a*, *socs3b*, *noxo1a, wnt4b, serpine1*, *stim1b*), “transmembrane signaling receptor activity” (*adora2b*, *ptger2a, lifrb, tlr5b, cxcr3.3, il4r.1, il4r.2, il6r*) and “signaling receptor activity” (*tnfrsf9a, tnfrsf18*), indicating activation of pro-inflammatory response (Figure 3C). Further, GO term linked to “chemotaxis”, “leukocyte migration” and “locomotion” were significantly enriched, alongside molecular function categories such as “chemokine activity” (*cxcl18b*, *cxcl11.6*, *cxcl11.1*, *cxcl8a*) and “CCR chemokine receptor” (*ccl38a.5, ccl35.1*, *ccl35.2, ccl34a.3*, *ccl34a.4 ccl39.6*) (Figure 3B, C and Table 6).

KEGG analysis revealed enrichment in 7 key inflammatory pathways, with the most prominent being the “c-type lectin receptor signaling” (*ptgs2a*, *bcl3, bcl10, irf1b*). Additional enriched pathways included “RIG-I-like receptor signaling” (*nfkbiaa, nfkbiab*), “NOD-like receptor signaling” (*tnfaip3, traf3*), “MAPK signaling” (*mknk1, mknk2a*), and “Toll-like receptor signaling” (*tlr5b, cd40*) (Figure 3D and Table 7). In short, at 2 hpA, macrophages enhance the transcription of genes clearly associated to immune cell activity, acute inflammation and defense mechanisms.

Among the top 50 down-regulated genes at 2 hpA, we identified several involved in the regulation of inflammation, including *tyk2*^41^*, sesn1*^42^ and *cd59*, a known inhibitor of complement-mediated cell lysis. Notably this list also includes the *cxcr4b* gene, which was shown to plays pleiotropic roles in wound repair and regeneration^43^ (Figure 3A and Table 5B).

These results reveal that macrophages first respond to the wound by adopting an acute pro-inflammatory program. While some genes were previously described in zebrafish activated wound macrophages, we identified novel genes underpinning early healing phase.

M1 macrophage signature in zebrafish is still poorly characterized and defining authentic markers for M1 could provide novel and improved tools to distinguish M1 macrophages in this specie. We hypothesized that the pro-inflammatory M1 signature would be conserved across various inflammatory conditions (sterile inflammation or bacterial infection). To test this, we compared the transcriptome of macrophages activated by a wound at 2 hpA with that of macrophages activated by *Salmonella enterica* infection at 4 hours post-infection, as we previously showed the pro-inflammatory profile of these macrophages^39^. We found 383 genes that were commonly up-regulated and 82 genes that were down-regulated in wound and infection (Table 8). The Spearman’s rho correlation coefficient of the log2 fold change was evaluated and showed that common DEGs are similarly modulated during early wound healing and early *Salmonella* infection (Figure 4). Thus, wound-activated macrophages at 2 hpA exhibited a pronounced M1-like core expression signature.

**Figure 4.**
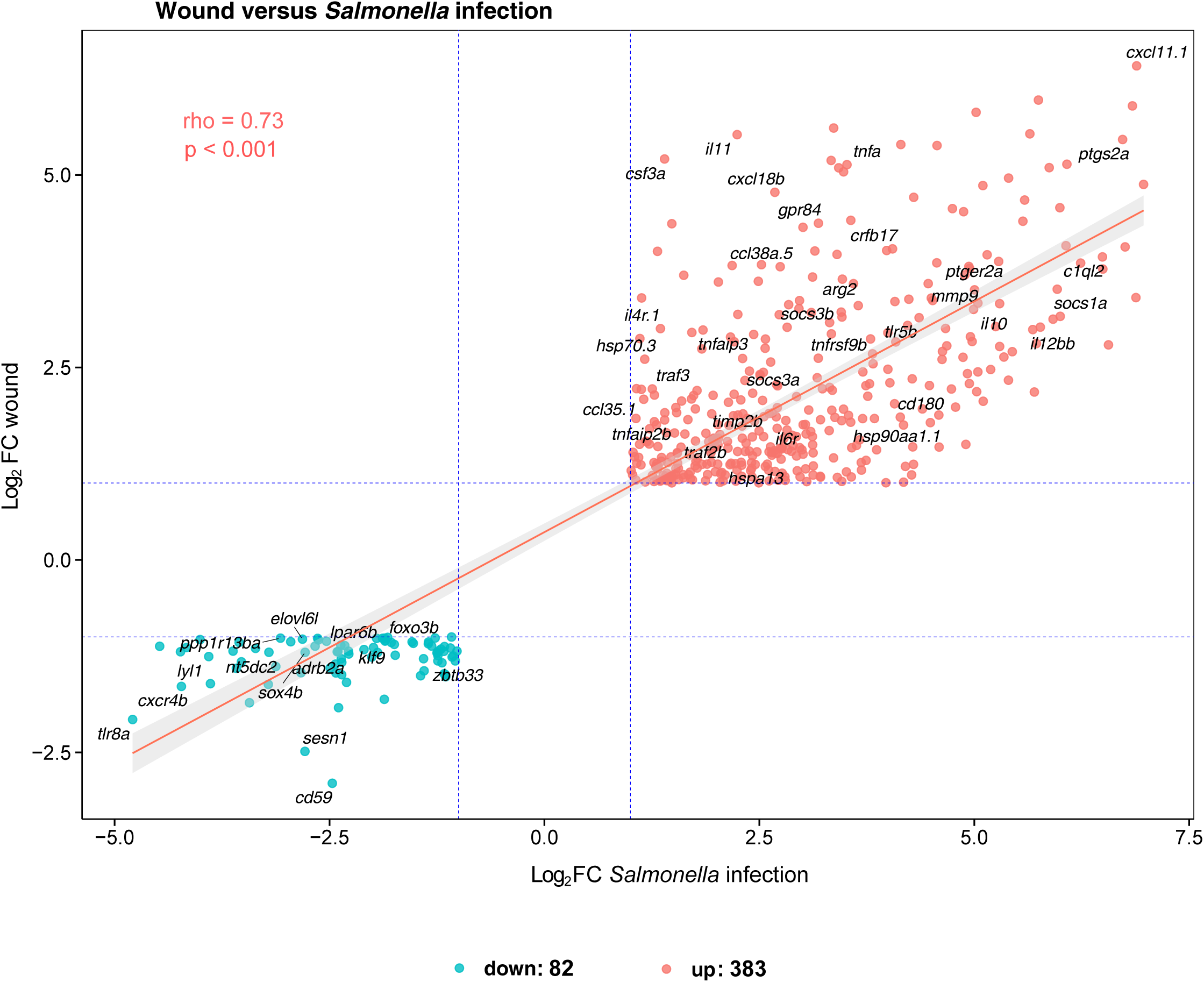
Correlation of gene expression fold changes between early wound healing and early *Salmonella* infection. The scatter plot displays the correlation of log₂ fold changes for genes that are commonly up- or down-regulated in both early wound healing and early *Salmonella* infection. Fold change correlation was calculated using Spearman’s correlation coefficient, based on 383 commonly up-regulated and 82 commonly down-regulated differentially expressed genes (DEGs). The y-axis represents the log₂ fold change in the early wound healing condition, while the x-axis shows the log₂ fold change during early *Salmonella* infection. The pink line indicates the linear regression fit. Dashed blue lines mark the fold change thresholds (log₂FC|1|) for both conditions. Each dot represents one gene, with color-coding indicating the regulation direction: pink for up-regulated and blue for down-regulated DEGs. Spearman’s correlation: rho = 0.73, *p* < 0.001.

### Five-hour-post-amputation macrophages display an immunoregulation and pro-reparative signature associated with glucose metabolism

Beyond pro-inflammatory macrophages, other wound-associated macrophage subtypes remain poorly characterized. We identified the *arginase 2* (*arg2*) gene, recently shown as an anti-inflammatory maker for macrophages and neutrophils during wound healing and mycobacteria infection^28^, was among the 50 up-regulated genes at 5 hpA. We also identified *itln1*, a lectin that reduces apoptosis and oxidative stress, previously described as anti-inflammatory^44^. Among the top up-regulated genes was *illr4*, a macrophage-inducible C-type lectin that promotes macrophage activation and inflammatory cell infiltration during sterile inflammation^45^ (Figure 5A and Table 9). Although not present in the top list, genes involved in lipid mediator class switching, such as *ptger2a and alox5ap* were also up-regulated.

**Figure 5.**
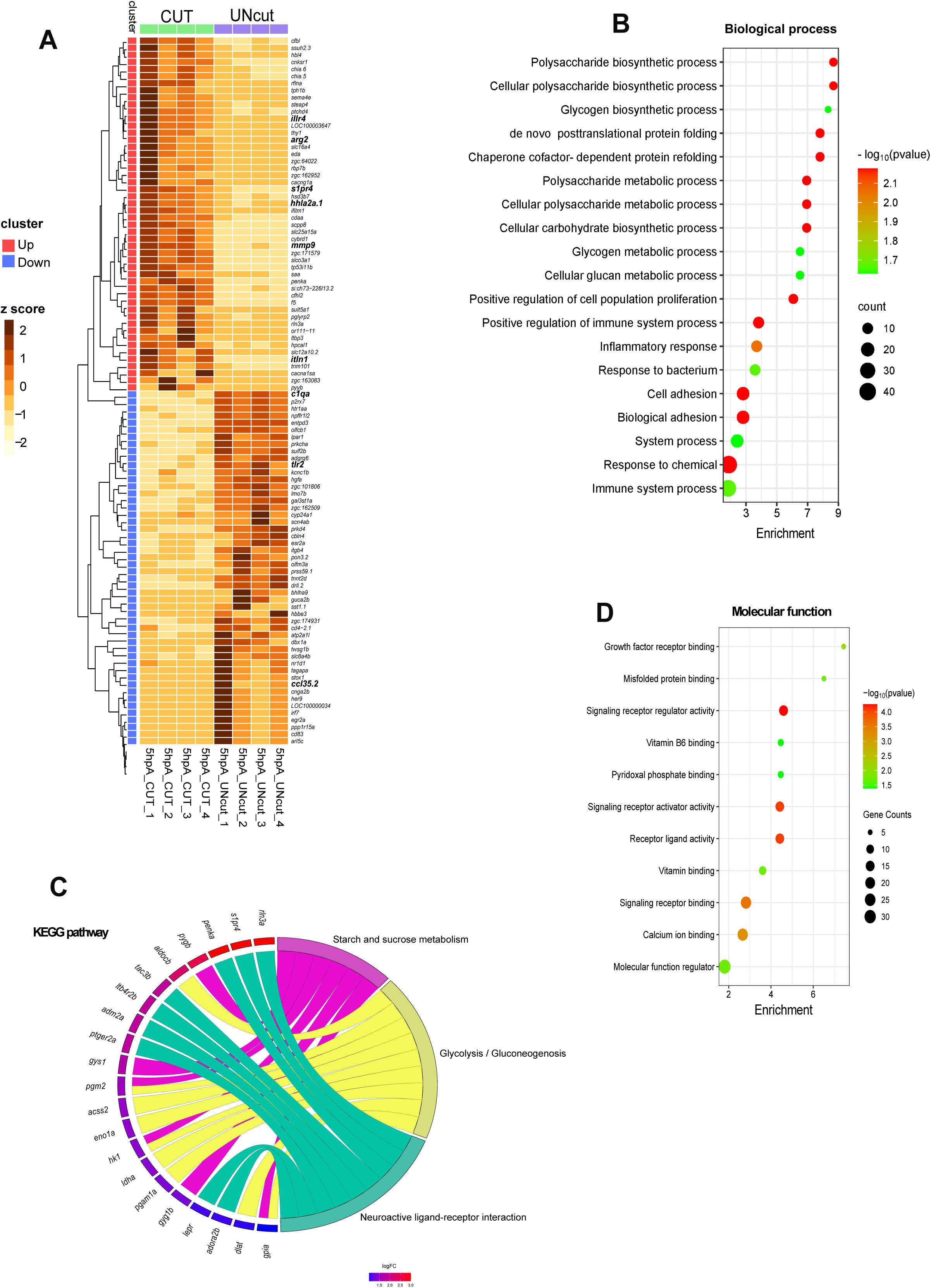
Transcriptome profiling of macrophages at 5 hpA in zebrafish larvae. (**A**) *Heat-map of differentially expressed genes.* The 50 most strongly up- and down-regulated genes are displayed. For each gene, normalized read counts were Z-score–transformed across all samples (columns = individual biological replicates). Colour scale: dark brown = high expression; light yellow = low expression. Clusters enriched in up-regulated genes are marked with red outlines; clusters enriched in down-regulated genes are marked in blue. Samples from amputated larvae (CUT) are labelled with green squares, and control samples (UNcut) with purple squares. Hierarchical clustering was performed with Ward’s minimum-variance method on log₂ fold-change values. **(B, D)** *Bubble plots of Gene Ontology enrichment.* Biological-process (**B**) and molecular-function (**D**) terms enriched among the up-regulated DEGs are shown. Dot size represents the number of genes in each term; dot colour encodes significance (FDR): red = higher significance (lower *p*-value), green = lower significance (higher *p*-value). The x-axis shows fold enrichment (gene ratio); the y-axis lists enriched GO terms. **(C)** *GO chord plot of Kyoto Encyclopedia of Genes and Genomes (KEGG)-associated terms.* Genes are connected to their associated pathways by ribbons. Genes are ordered by log₂ fold change (red = strong up-regulation; blue = strong down-regulation).

Overrepresentation analysis was carried out on the DEGs using gene ontology (GO) and the KEGG pathways analysis. GO analysis at 5 hpA showed little overlap with processes detected at 2 hpA. At this later stage, up-regulated genes were mapped in glucose metabolism-related categories: “polysaccharide biosynthetic / metabolic process”, “glycogen biosynthetic/metabolic process” and “cellular carbohydrate biosynthetic process” (Figure 5B). KEGG pathway enrichment further confirmed a shift toward metabolic pathways, particularly: “glycolysis” (*aldocb, acss2, eno1a, hk1, idha, pgam1a*, *dlat*, *gpia*) and “starch and sucrose metabolism” including glycogenesis (*gys1*, *pgm2*) and glycogenolysis (*pygb* and *gyg1b)* (Figure 5C and Table 10). These results suggest that 5hpA-macrophages rely primarily on glucose-fueled glycolysis and the glycogenesis-glycogenolysis cycle to meet energy demand and support their functional activities.

Among molecular functions, up-regulated DEGs were enriched GO terms “signaling receptor activity” (*tnfsf13b*, *gpia*, *pyyb*, *adm2a*, *rln3a*, *eda*, and *ccl34a.3*) (Figure 5D). This category also featured the expression of key homeostatic signaling regulators such as interleukins *il6*, *il10* and *penka*^46^. Of particular interest in the context of tissue repair, we identified up-regulated genes associated with the GO term “positive regulation of cell population proliferation,” including growth factors such as *fgf6a, hbegfa, hbegfb,* and *vegfab* (Figure 5B, D). These factors, recognized as pro-resolving markers, have been implicated in initiating tissue repair by promoting fibroblast and keratinocyte proliferation, supporting vasculogenesis, and epithelialization^1^. In addition, enriched GO terms linked to “chaperone cofactor-dependent protein refolding” and “de novo posttranslational protein folding” were enriched, highlighting molecular chaperones involved in protein quality control during tissue repair (Figure 5B, D). Notably, several members of the heat shock protein family, including *hspa9, hspa8b*, *hsp70.1 and hsp70.2*^47,48^, were among the up-regulated genes. Interestingly, genes known to be involved in wound healing, such as *annexin* (*anxa2a*, *anxa1a*, *anxa6*, *anxa13*)^49^, were identified within the enriched GO term "calcium ion binding". (Figure 5D and Table 11).

In addition, the KEGG analysis revealed the “Neuroactive ligand-receptor interaction” pathway, which encode GPCRs or ligands that bind to GPCRs. Some of these genes are involved in immune response modulation (*adora2b*, *penka*, *ptger2a*, *adm2a*, *ltb4r2b*, *lepr* and *s1pr4*) (Figure 5C and Table 10).

By contrast, GO analysis of down-regulated DEGs at 5 hpA revealed enrichment in terms related to immune modulation, including “regulation of immune response,” “regulation of defense response,” and “positive regulation of the MAPK cascade” (Table 12). It is noteworthy that many genes associated with the acute pro-inflammatory phase (2hpA) presented in Table 2A are not differentially expressed at 5 hpA (Table 3A). Genes such as *il1b*, *il12bb*, *tnfa*, *cxcl8a*, *cxcl18b*, *ccl34a.4*, *ccl38a.5*, and *traf2b*, which were up-regulated at 2hpA, showed no significant changes at 5 hpA. These findings suggest that by this time point, macrophages begin to downregulate genes associated with the acute inflammatory response, indicating a transition toward the resolution phase and progression of wound healing.

Overall, these findings suggest that by 5 hpA, macrophages undergo a metabolic shift towards glucose and glycogen metabolism while regulating the acute pro-inflammatory phase and transcribing genes to promote repair.

### Some DEGs are shared between times 2 hpA and 5 hpA

Several differentially expressed genes (DEGs) were common between 2 hpA and 5 hpA (Figure 1D). We analyzed the correlation of log2 fold change between all commonly regulated genes at 2 and 5 hpA using the Spearman log2 fold change correlation coefficient (Figure 6). We found an acceptable relationship between the log2 fold change of both conditions (2 hpA and 5 hpA) (rho = 0.7 with a p-value < 0.001), indicating that a part of the genes display similar modulations during first steps of wound healing. However, some genes subset exhibiting rapid modulation during early wound healing. In particular, 14 DEGs were up-regulated at 2 hpA and subsequently down-regulated by 5 hpA (Figure 1D and Table 13A). These genes encode key regulators of the host immune response that induce pro-inflammatory cytokines and interferon transcription. For instance, *nfkb2*, *nfkbib*, and *zfand5a* serve as central activators of pro-inflammatory response, while signaling factors such as *traf3*, *fosl1a*, *mknk2a*, *tagapa*, and *irf1b* further modulate this response. Additionally, chemokine *ccl35.2* and the receptor *crfb15* were identified, suggesting that the pro-inflammatory pathway is rapidly down-regulated to optimize healing and minimize tissue damage.

**Figure 6.**
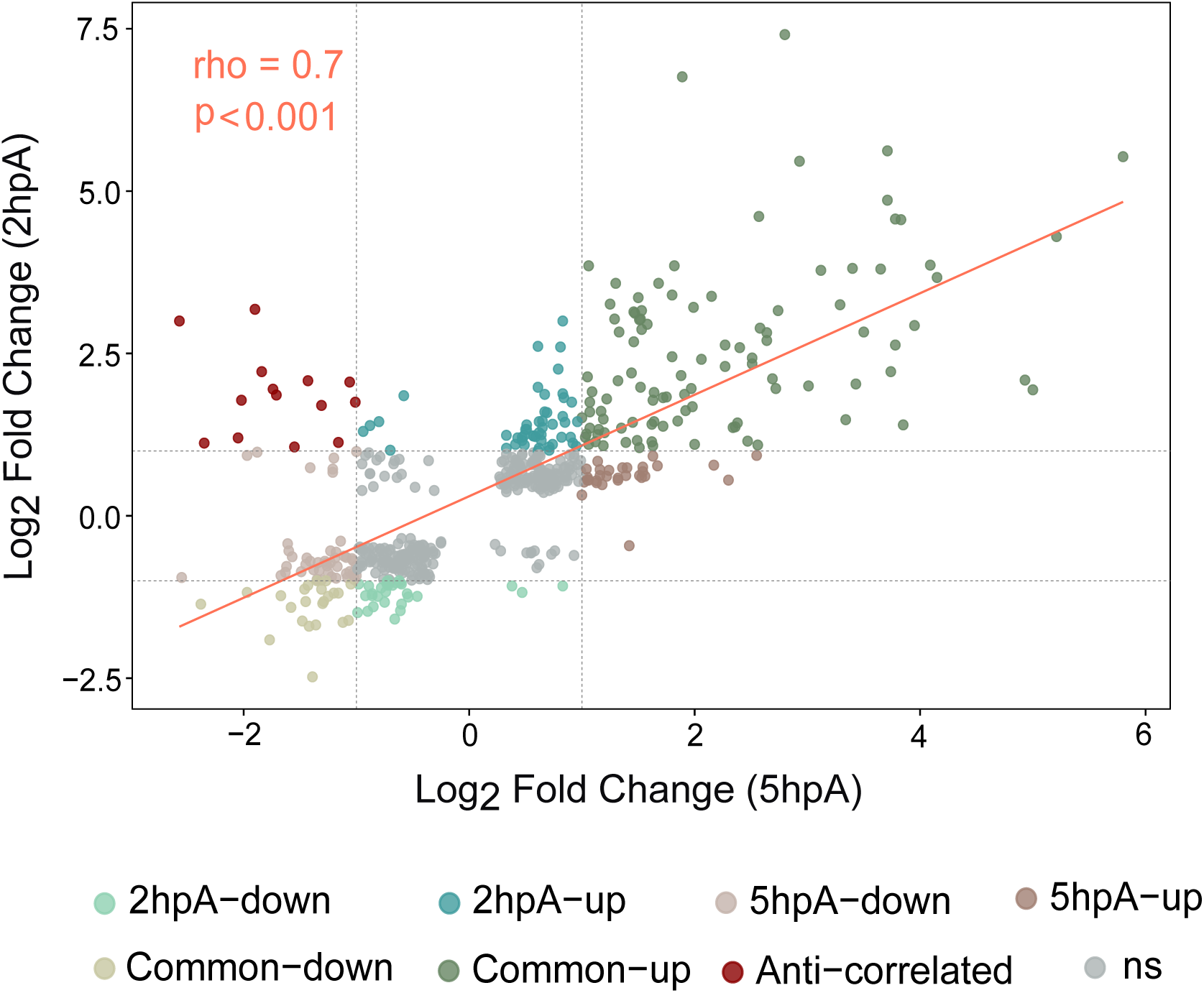
Correlation of gene expression fold changes between 2hpA and 5hpA. The scatter plot depicts the correlation of log₂ fold changes in gene expression between the 2 hpA and 5 hpA conditions for the 141 shared DEGs. Each point represents a gene. The x-axis shows the log₂ fold change at 5 hpA, and the y-axis shows the log₂ fold change at 2 hpA. The dashed vertical and horizontal lines indicate the fold change threshold (log2FC |1|) for both time points. The color-coding differentiates up- and down-regulated genes unique to each time point (shades of green for 2 hpA, shades of brown for 5 hpA), commonly regulated genes, anti-correlated genes (dark red), and non-significant (ns) genes (grey). The pink line represents the linear regression. Fold change correlation was assessed using Spearman’s correlation coefficient (rho = 0.7, p < 0.001).

Of the differentially expressed genes (DEGs), 104 up-regulated at 2 hpA remained elevated at 5 hpA (Figure 1D). Notably, several showed a nearly fivefold increase at 5 hpA (Table 13B). They include genes are involved in mitochondrial oxidative metabolism such as, *ncf2* and *cybrd1*, underlying the mitochondria’s key role in wound healing. Genes like, *s1pr4* (immunoregulation), *prdx1* (oxidative stress protection), and *dgkaa*, associated with immunoregulatory functions in macrophages^50^, showed enhanced expression at 5 hpA. Similarly, *cdaa*, linked to the pyrimidine salvage pathway essential for nucleotide balance^51^ and *slc25a15a, a* mitochondrial carrier involved in urea metabolism and nitric oxide clearance^52^ exhibited ∼7-fold increases. Moreover, genes coding for wound healing effectors (*timp2b*, *arg2*, and *tp53i11b)*, and enzymes remodeling extracellular matrix exhibited an average 1.5-fold increase at 5 hpA (Table 13B).

Taken together, this finding suggests that in wound-macrophages, the early activation of healing mechanisms during acute inflammation phase sets the stage for subsequent resolution and tissue repair.

### Macrophage reprogramming becomes attenuated during the later stages of healing

By contrast to earlier stages, at 29 hpA, limited number of DEGs (N=45) were identified (Table 4A). Among them, 28 were found up-regulated. Seven genes remained consistently regulated across all three time points (2 hpA, 5 hpA, and 29 hpA), albeit with variable fold changes (Table 13C). They included genes encoding extracellular matrix remodeling factors (*mmp9* and *mmp13*), which play critical roles in maintaining tissue homeostasis and facilitating epithelial repair during wound healing. The sustained expression of these genes suggests a prolonged role in wound repair and may represent a general signature of wound-associated macrophages.

Among the up-regulated genes at 29hpA time point, we identified genes associated with immune regulation (pdlim1, chia.6, padi2) and host defense (*npsn, nfkb2, mpx, cp*). Further several up-regulated genes were involved in intracellular trafficking (*bicdl2l, tom1l2, snx11, sacm1la*), cytoskeletal dynamics (*scinlb, bcas3, arhgap1*) and autophagy/mitophagy (*prkcda, ulk1a, rab7b, fnip1*) (Table 4A), suggesting that 29-hpA macrophages display functions relying on membrane trafficking and degradative pathways such as efferocytosis and fine-tuning signaling pathways by recycling of surface receptors. Further increased levels of mitophagy related genes suggest a metabolic reprogramming linked to mitochondria. Importantly at 29 hpA, 22 genes (9 up-regulated and 13 down-regulated genes) were unique to this time point, representing a signature of macrophage associated to late healing (Table 4B, C)

This finding demonstrates that by 29 hours post-injury, the reprogramming of macrophages is deeply attenuated. It also reveals a distinct wound-healing signature marked by the non-inflammation, the matrix remodelling and membrane trafficking, supportive of tissue repair.

## 4. Discussion

The full transcriptome of zebrafish macrophages associated to wound healing has not been mapped in detail. Here, we report the first comprehensive analysis of transcriptional profiles of macrophages during wound healing in zebrafish. Bulk RNA sequencing of purified zebrafish macrophages at different times following fin fold injury allowed us to characterize the highly dynamic changes in gene expression occurring in macrophage reflecting shifts in phenotype over the first hours.

Specifically, we show that 2 hpA-macrophages display a strong pro-inflammatory phenotype characterized by the expression of genes involved in immune cell activation, acute inflammation and defense response. Indeed, based on their mRNA profile, 2 hpA-macrophages may secrete variety of chemokines and inflammatory cytokines, such as *il-1b, il-12bb*, *il-6* and *tnfa,* respond to them by using chemotaxis and by robustly activating inflammatory signaling pathways. Key members of pro-inflammatory pathways including *nfkbia/b*, *nfkb2, tlr5b* and *traf3* are upregulated. In mammals, TRAF3 plays different roles in the inflammatory signaling pathways mediated by TNFR, TLR, and RLR by participating in different protein complexes, including the direct activation IKK complex in NF-κB signaling pathway^53^ and actuation of MAPK pathway. All these inflammatory pathways were found enriched in the zebrafish 2 hpA-macrophages. We detected expression of *ptgs2a* (also known as *cox2*), a key enzyme in eicosanoid synthesis. In addition, the chemokine *cxcl11.1* and its receptor *cxcr3.3* were identified. *Cxcl11* has been reported as a macrophage polarization marker during mycobacterial infection^23^ while *cxcr3.3* regulates macrophage trafficking by antagonizing the *cxcl11*/*cxcr3.2* axis^54^. Consistent with previous findings, *cxcl8a*, *cxcl18b*, and the receptor *ccl38a.5* are up-regulated in 2 hpA macrophages; these chemokines are known to induce neutrophil chemotaxis in zebrafish^55, 56^. While 2 hpA-macrophages upregulate pro-inflammatory genes early in response to injury, several genes that inhibit these pathways are also induced (e.g., *il10*, *arg2*), most likely to exert negative feedback and limit excessive inflammation. Finally, while we confirmed conserved pro-inflammatory markers in M1-like zebrafish macrophages^57, 58^, our findings also reveal a broader set of markers, signaling pathways, and effectors.

In this study, we show that 5-hpA-macrophages downregulate many of the inflammatory genes elevated at 2hpA, including *nfkb2*, *nfkbib*, *zfand5a*, *traf3*, *fosl1a*, *mknk2a* and *irf1b*. Concomitantly, they strongly up-regulate genes modulating inflammation such as *adora2b* and *lepr* in line with a transition period from acute to resolution-phase. In addition, 5 hpA-macrophages exhibit expression of growth factor like *fgf6a, hbegfa/b and vegfab* and activation of cell proliferation. While Fibroblast growth factor (Fgf) signaling is essential for fin regeneration in zebrafish, the specific role of *fgf6a* has not yet been studied in this specie^59^. In mammals, FGF6 enhance muscle regeneration^60, 61^. Further *hbegf* and *vegf* family members are pro-regenerative macrophage makers, however their exact role in fish fin fold healing is unknown. Five-hpA-macrophages also express key genes involved in the production of lipid class switch and pro-resolving lipid mediators such as *alox5ap, alox12*, *alox5a*. These findings align with zebrafish studies showing that Alox12 upregulation during wound healing promotes anti-inflammatory lipid signaling^62^. Importantly these macrophages show phenotypic shifts at the transcriptomic level that are associated with tissue repair, highlighted by expression of annexins which are Ca^2+^-regulated membrane binding proteins responsive to cellular stress. This suggests that 5 hpA macrophages are transitioning from an inflammatory to a reparative state, displaying a mixed phenotype with features of both M1- and M2-like macrophages.

At 29 hpA, macrophages exhibit a non-inflammatory, reparative phenotype. They markedly up-regulate extracellular-matrix–remodeling genes, notably *mmp9* and *mmp13a*, metalloproteinases previously shown to be essential for zebrafish regeneration^63,64^. Comparable ECM remodelling-focused programs have been described in other injury models including zebrafish fin repair and murine and human cardiac injury, where macrophages shape scar formation and matrix composition^65,66^. Beyond matrix genes, 29 hpA macrophages increase transcripts linked to actin remodeling, intracellular trafficking, and autophagy, consistent with active efferocytosis and receptor recycling. The relatively small set of differentially expressed genes at this late stage suggests that major transcriptional reprogramming has subsided, reflecting a transition toward stabilization rather than continued plasticity.

Interestingly the discovery of immunometabolic features at 5 hpA, provides insights into macrophage functional metabolic reprogramming during wound healing in zebrafish. Macrophage metabolism plays a key role in determining polarization states in mammals^67, 68^. Pro-inflammatory macrophages are typically driven by glycolysis in response to bacterial and inflammatory stimuli, whereas anti-inflammatory (M2-like) macrophages preferentially utilize oxidative phosphorylation (OXPHOS) and the TCA cycle. It is widely accepted that metabolic reprogramming precedes and supports the resolution of inflammation^67, 68^. Strikingly, our data reveal a metabolic shift in macrophages at 5 hpA, characterized by the upregulation of genes involved in glycolysis, glycogenolysis, gluconeogenesis, and activation of the pentose phosphate pathway (PPP). Although transcripts associated with the TCA cycle were present, we initially observed no strong enrichment of genes related to fatty acid oxidation or oxidative phosphorylation (OXPHOS). Given this unexpected result, we re-analyzed the up-regulated DEGs with moderate fold changes (log2FC |0|), and identified several genes linked to OXPHOS (including ndufs5, uqcrfs1, ndufv2, ndufa10, atp6v1ba, atp5pf, uqcrc2a, ndufab1b, ndufs3, cox5aa, cox6a1, cox6b1, cox6b2, cox7c, cox7b, cox10, cox4i1, cox11, cox17, atp5mc3a, ndufb4, atp5f1d, atp6v0a1a, atp6v1c1a, ndufs1, ndufb10, uqcrq, sdhdb, ppa1b). This suggests that while glucose metabolism is the predominant energy source at this stage, a modest OXPHOS signature is also emerging. These findings highlight the metabolic complexity of macrophages *in vivo*, where multiple pathways likely interact and compensate to support functional demands.

This metabolic profile contrasts with that observed during skeletal muscle regeneration in mammals, where macrophages progressively shift toward TCA cycle activity as they transition from pro- to anti-inflammatory states^69^. Similarly, studies using myocardial infarction (MI) models have shown that macrophages undergo a metabolic shift toward oxidative phosphorylation during the post-MI remodeling phase^70^. In contrast, our findings align with previous zebrafish studies reporting increased glycolysis in macrophages at 24 hpA, a process shown to be essential for regenerative function^71^. This suggests that during the transition phase in zebrafish, macrophages may rely on glucose metabolism to support their function.

Interestingly, 5-hpA-macrophages display enrichment in gene of glycogen metabolism. Glycogen, a glucose-based polymer, functions as a key carbohydrate reserve within cells, readily mobilized to meet energy demands. Glycogen synthase (*gys1*) and glycogen phosphorylase (*pygb*) that are respectively involved in glycogenesis and glycogenolysis, were found upregulated at this stage, suggesting that macrophages specifically engage in glycogen cycling for energy production and functional activity. The presence of glycogen stores in human and rodent myeloid cells was reported several decades ago^72^. A recent study showed that in mice, M1 macrophages actively synthesize and degrade glycogen^73^. Similarly, human M1 and tumor-associated macrophages store glycogen and perform gluconeogenesis. This glycogen-centered metabolic program may support cytokine production and phagocytosis under low-glucose conditions. Further functional investigations are needed to determine whether zebrafish macrophages adapt to physiological stress through metabolic specialization, potentially by cycling between gluconeogenesis and glycogenesis, to maintain energy homeostasis.

One intriguing question is when appear the resolutive macrophages? Interestingly, we observed that genes associated with tissue repair and resolution, such as *timp2b*, begin to be expressed as early as 2 hours post-injury. This suggests that a subset of macrophages with reparative or pro-resolving functions may be activated very early, in parallel with those driving acute inflammation. As we previously demonstrated, macrophages at the wound site include both *tnf-a.a*–positive and –negative subsets, suggesting the coexistence of functionally distinct populations^8^. Rather than a strictly sequential transition from pro-inflammatory to anti-inflammatory states, our data support a model in which heterogeneous macrophage populations with distinct functional programs are recruited or activated simultaneously.

Our study demonstrates that zebrafish macrophages exhibit dynamic phenotypic plasticity in response to the changing microenvironment of a wound. These phenotypes are not static but are tightly regulated over time, with the majority of transcriptional reprogramming occurring within a relatively short window and showing marked attenuation by 29 hpA. While previous transcriptomic studies in zebrafish focusing on macrophage program during injury^74^, did not fully capture the pro-inflammatory phase of macrophage activation, our study provides the first comprehensive description of both the pro-inflammatory response and the onset of resolution.

Additional transcriptomic and metabolic analyses at intermediate time points between 6 and 29 hpA will be essential to uncover potential metabolic transitions and signaling pathways involved in intermediate stages of wound healing. It is worth noting that our macrophage isolation protocol, which employed enzyme-free dissociation and FACS purification, was designed to minimize artificial activation under steady-state conditions^29^. Although few transcriptional changes were observed at 29 hpA, it remains likely that macrophages exhibit altered protein expression at this stage that are not captured by bulk transcriptomic, underscoring the importance of complementary proteomic approaches in future studies. Additionally, the use of bulk RNA sequencing, while offering a comprehensive overview of gene expression, inherently masks cellular heterogeneity. Beside these limitations, we were able to identify bona fide makers for polarized macrophages across wound healing in zebrafish. Finally, we note that data from various injury paradigms, organs and species suggest that the molecular pathways discovered in this study are conserved across multiple biological contexts.

## Acknowledgements

This work was undertaken with support from the fish facility of ZEFIX-LPHI (University of Montpellier), Catherine Gonzalez (LPHI, CNRS) and the MGX-Montpellier GenomiX platform and the qPCR Haut Debit platform of the University of Montpellier. We are grateful to Georges Lutfalla for his experimental help and secure funding. MGX acknowledges financial support from France Génomique National infrastructure, funded as part of “Investissement d’Avenir” program managed by Agence Nationale pour la Recherche (contract ANR-10-INBS-09). We acknowledge the imaging facility BioCampus Montpellier Ressources Imagerie (MRI), member of the national infrastructure France-BioImaging supported by the French National Research Agency (ANR-10-INSB-04, “Investments for the future”).

## Authorship contributions

C.B-P, M.N-C. performed and conceived zebrafish experiments. C.B-P performed RNA quality validation. R.O. performed zebrafish experiments. S.B. performed FACS experiments. S.G. performed library preparation, RNA-sequencing and validations. A.L. performed bioinformatics and statistical analysis. L.M. advices for RNA-Seq, performed data processing, analysis and figures. M.N-C performed supervision and secured funding. C.B-P and M.N-C wrote the manuscript. All authors reviewed the manuscript and approved the submitted version.

